# Soluble pathogenic tau transmission to astrocytes drives cellular senescence and neurovascular uncoupling in a model of Alzheimer’s tauopathy

**DOI:** 10.64898/2026.09.08.750215

**Authors:** Haneen Makhlouf, Angela O Dorigatti, Stephen Hernandez, Nicholas de Rosa, Lauren Miller, Stacy Hussong, Deborah Holstein, James Lechleiter, Sreemathi Logan, Rakez Kayed, Veronica Galvan

**Affiliations:** Department of Neurosurgery, University of Oklahoma Health Sciences Center, Oklahoma City, OK, USA.; Oklahoma City Veterans Health Care System, 921 NE 13th Street, Oklahoma City, OK, 73104, USA; Illumina, Inc. 5200 Illumina Way, San Diego, 92122, USA; Department of Molecular Medicine, University of Texas Health San Antonio, San Antonio, TX, USA; Department of Cellular & Structural Biology, South Texas Research Facility Neuroscience Center, University of Texas Health Science Center at San Antonio, San Antonio, TX, USA; Department of Biochemistry and Physiology, University of Oklahoma Health Sciences Center, 940 Stanton L Young Blvd, Oklahoma City, OK, 73104, USA; Center for Geroscience and Healthy Brain Aging, University of Oklahoma Health Sciences Center, 940 Stanton L Young Blvd, Oklahoma City, OK, 73104, USA; Departments of Neurology, Neuroscience and Cell Biology, University of Texas Medical Branch at Galveston, 301 University Blvd, Galveston, TX, 77555, USA; Mitchell Center for Neurodegenerative Disease, University of Texas Medical Branch at Galveston, 301 University Blvd, Galveston, TX, 77555, USA; Sealy Center for Vaccine Development, University of Texas Medical Branch at Galveston, 301 University Blvd, Galveston, TX, 77555, USA

## Abstract

Pathogenic tau oligomers are transmitted between neurons and non-neuronal cells and may contribute to cellular dysfunction in Alzheimer’s disease (AD). We previously demonstrated that soluble tau aggregates accumulate in brain microvascular endothelial cells, inducing cellular senescence and microvascular dysfunction in a mouse model of tauopathy. Here, we show that soluble pathogenic tau is also transmitted to astrocytes, where it induces mitochondrial oxidative stress and a senescence-associated phenotype (SASP). Single-cell RNA sequencing of hTau mouse cortex identified astrocytes among the most transcriptionally altered cell types, with reduced expression of electron transport chain genes and increased expression of stress and inflammatory markers. This transcriptional pattern was also observed in hTau brain and astrocyte-enriched fractions. Soluble tau aggregates transmitted to primary human astrocytes through a heparin-sensitive process and rapidly decreased ATP levels and increased mitochondrial ROS. These mitochondrial changes preceded detectable microtubule destabilization and were followed by induction of SASP and cell-cycle-arrest-associated markers. Scavenging mitochondrial ROS with MitoTEMPO reduced tau-induced SASP cytokine and senescence-associated responses in vitro and in vivo, supporting a causal contribution of mitochondrial oxidative stress to senescence induction. Neurons cocultured with astrocytes undergoing tau-induced senescence exhibited reduced dendritic spine density, branching, and dendritic area, demonstrating non-cell-autonomous effects on neuronal structure. Astrocyte-targeted SOD2 overexpression ameliorated deficits in evoked cerebral blood-flow responses in hTau mice. Together, these findings identify mitochondrial oxidative stress as a mechanistic link between pathogenic tau transmission and astrocyte senescence-associated dysfunction and implicate impaired mitochondrial antioxidant defense in neuronal and neurovascular abnormalities associated with tauopathy.

## Introduction

Tau is a microtubule-associated protein that, under normal conditions, stabilizes neuronal microtubules^1^. In Alzheimer’s disease (AD) and primary tauopathies, tau becomes abnormally phosphorylated, dissociates from microtubules, misfolds, and assembles into higher-order structures^2,3^. Tau oligomers are soluble assemblies that can precede the formation of mature fibrillar tau aggregates, including insoluble neurofibrillary tangles^4,5^. These oligomeric species are neurotoxic, and their intracerebral administration into wild-type mice is sufficient to induce synaptic and memory deficits^5^. These findings suggest that tau oligomers can act as pathogenic species rather than simply as intermediates in tangle formation. Tau oligomers can be detected using conformation-specific antibodies that recognize oligomeric tau but not monomeric or fibrillar forms^6^.

Neuronal activity promotes extracellular tau release^7,8^ and enhances tau propagation through anatomically connected brain regions^9,10^. Misfolded, oligomeric tau can be taken up by recipient neurons, where it promotes the misfolding and aggregation of endogenous tau, supporting a prion-like model of tau propagation^11–13^. Macropinocytosis is one route of tau uptake and can depend on tau binding to cell-surface heparan sulfate proteoglycans (HSPGs)^14^. Low-density lipoprotein receptor-related protein 1 (LRP1) also mediates neuronal tau uptake and spread, indicating that multiple cell-surface mechanisms contribute to tau entry^15^. Together, these findings support extracellular tau transmission as an important mechanism through which tau pathology can propagate between cells and across connected brain regions during AD progression.

While the prion-like propagation model largely focused on neurons, astrocytes can also internalize extracellular tau. Astrocytes contact many synapses and extend end-feet along cerebral blood vessels^16–18^, positioning them to encounter extracellular tau and potentially influence both neuronal and vascular function. Tau pathology is a characteristic feature of primary tauopathies and is increasingly detected in the aging human brain^19,20^. Transmission of pathogenic tau to different cell types may have distinct functional consequences. Transcriptomic studies in AD mouse models and the aging human brain show that disease-associated astrocyte states emerge early and expand as disease progresses^21–23^. Astrocytes’ transcriptional responses vary across brain regions and stages of pathology, from normal aging to AD. In primary astrocytes and mouse models, increased expression of the transcription factor TFEB increases uptake and degradation of extracellular tau, reducing tau spread^24^. Human astrocytes derived from induced pluripotent stem cells and exposed to tau fibrils from AD brains show changes in their ability to clear tau and in their tau-seeding activity, along with transcriptional changes that overlap with reactive-astrocyte patterns seen in AD patient brains^25,26^.

Astrocytes clear synaptic glutamate and translate neuronal activity into vasodilatory signals that contribute to neurovascular coupling^27–31^, functions that impose substantial metabolic demands on astrocytes^32–35^. Neurovascular deficits are among the earliest abnormalities detected in AD patients^36,37^, yet the impact of tau transmission to astrocytes on cellular and neurovascular function remains poorly understood. During aging, cells accumulate damage beyond repair capacity^38^; stress can promote senescence, a state characterized by stable cell-cycle arrest and development of a senescence-associated secretory phenotype (SASP)^39–44^. Senescent glial cells accumulate in tauopathy models^45,46^, and eliminating senescent glial cells prevents tau-dependent pathology and cognitive decline in a tauopathy model^47^. More broadly, clearance of p16-positive cells delays age-associated disorders^48^, while p16-expressing microglia and endothelial cells promote tauopathy-associated neurovascular abnormalities^49^. In another tauopathy model, telomere-driven senescence accelerates tau pathology, neuroinflammation, and neurodegeneration^50^. Mitochondrial dysfunction is an established inducer of senescence^51,52^, and reciprocal signaling between reactive oxygen species (ROS) and cell cycle regulators such as p21 can reinforce SASP and maintain senescence-associated arrest^53^. Prior work showed that soluble pathogenic tau enters brain microvascular endothelial cells and causes senescence and microvascular dysfunction^54^. Senescent astrocytes can also alter signaling to brain endothelial cells^55^. Whether tau transmission to astrocytes disrupts mitochondrial function and triggers senescence remains poorly understood.

This study examines how pathogenic tau transmission to astrocytes affects mitochondrial function, cellular senescence, neuronal structure, and neurovascular coupling. The central hypothesis is that tau-induced mitochondrial oxidative stress promotes astrocyte senescence and disrupts synaptic integrity and neurovascular coupling. We used single-cell RNA-seq of cortical tissue to characterize transcriptional changes in astrocytes. We assessed the impact of soluble tau oligomers on primary human astrocyte microtubule stability, ATP production, mitochondrial reactive oxygen species (ROS), and the timing of senescence-associated secretory phenotype (SASP) and cell-cycle-arrest marker induction. To determine the causal role of mitochondrial ROS in astrocyte senescence, we used the mitochondria-targeted antioxidant MitoTEMPO. We evaluated the influence of tau-induced senescent astrocytes on neuronal structure by co-culturing them with primary neurons and measuring dendritic spine density, dendritic branching, and SASP factor contribution. Finally, we investigated whether astrocyte-targeted SOD2 overexpression could enhance evoked cerebral blood flow responses in hTau mice in vivo.

## Results

### Soluble tau aggregates enter primary human astrocytes via a heparin-sensitive mechanism and induce a senescent, pro-inflammatory phenotype

To investigate the effects of soluble pathogenic tau on astrocytes, primary human astrocytes were incubated for 48 hours with 40 µg/mL soluble oligomeric tau (O.Tau). Because cellular uptake of pathogenic tau may depend on heparan sulfate proteoglycans^14^, additional groups were treated with O. Tau in the presence of 0.5 mg/mL heparin sodium. Heparin alone and KRT8, a control protein with similar molecular weight and isoelectric point to tau, served as controls. Confocal microscopy revealed tau-positive puncta in O. Tau-treated astrocytes, distributed throughout the cytoplasm and along the cytoskeletal network (**Fig. 1A**). In astrocytes treated with both heparin and tau, intracellular tau staining was reduced, indicating that the uptake mechanism for tau aggregates is sensitive to heparin (**Fig. 1A**). Exposure to O. Tau increased cell area without significantly altering nuclear area (**Fig. 1B–D**).

**Figure 1:**
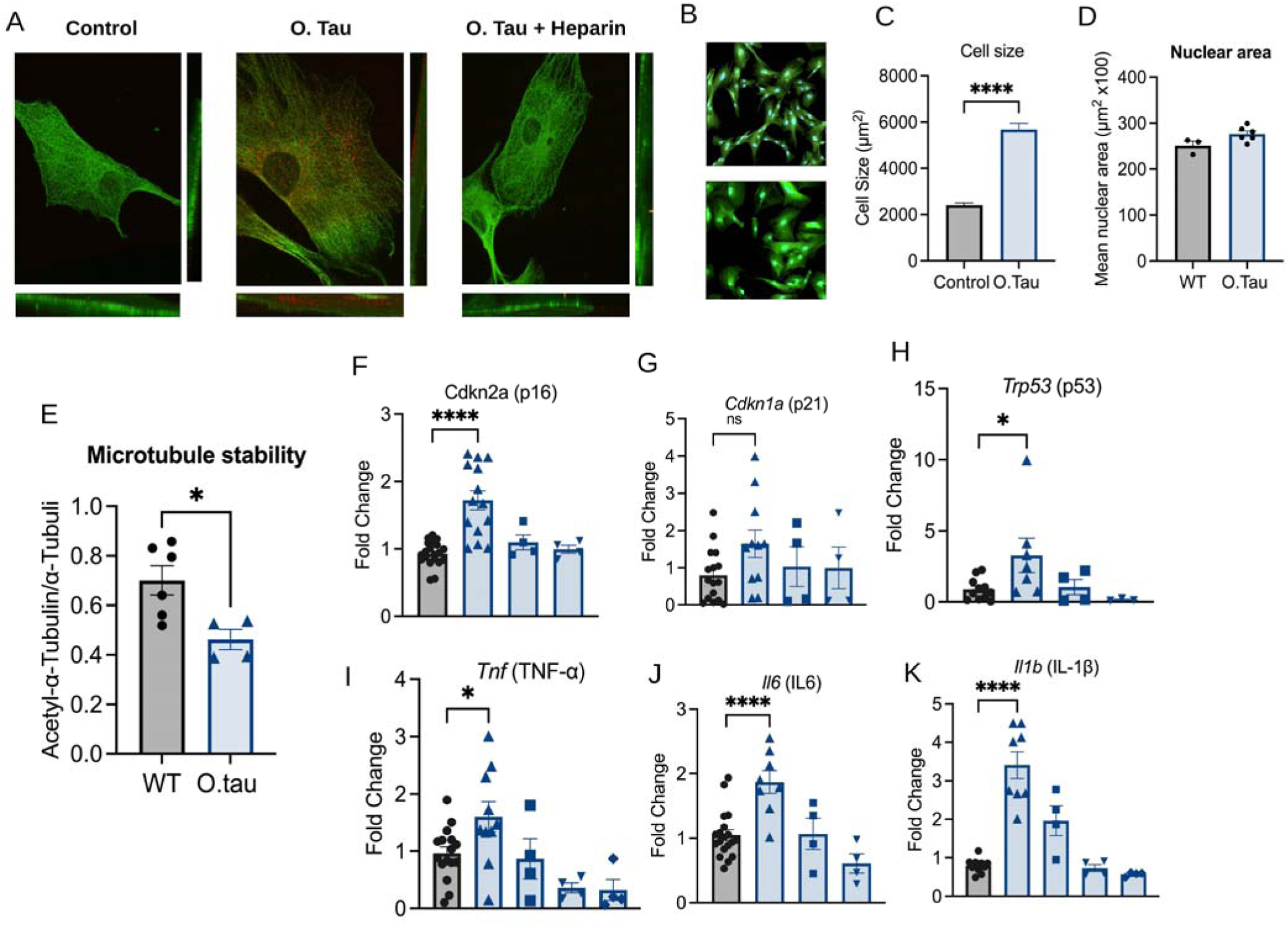
Soluble tau aggregates enter primary human astrocytes via a heparin-sensitive mechanism and induce a senescent, pro-inflammatory phenotype. (**A**) Representative confocal images of primary human astrocytes treated for 48 hours with vehicle, soluble oligomeric tau (O. Tau; 40 µg/mL), or O. Tau plus heparin sodium (0.5 mg/mL). Tau-positive puncta are shown in red and the astrocyte cytoskeleton in green. Orthogonal projections show intracellular tau localization.(**B**) Representative images of astrocyte morphology after vehicle or O. Tau treatment.(**C, D**) Quantification of astrocyte cell area and nuclear area, respectively.(**E**) Microtubule stability assessed as the ratio of acetylated a-tubulin to total a-tubulin.(**F-H**) Relative expression of the senescence-associated genes CDKN2A (p16), CDKN1A (p21), and TP53 (p53) respectively.. (**I–K**) Relative expression of the inflammatory genes TNF, IL6, and IL1B respectively. Heparin alone and KRT8, a control protein with a molecular weight and isoelectric point comparable to tau, were included as controls. Data are presented as mean ± SEM from five independent experiments. Each symbol represents an individual measurement. Statistical tests are specified in the Methods. *P < 0.05; ***P < 0.0001; ns, not significant.

Treatment with O. Tau decreased the ratio of acetylated α-tubulin to total α-tubulin, indicating reduced microtubule stability after 48 hours (**Fig. 1E**). O. Tau exposure also upregulated the senescence-associated genes CDKN2A and TP53, while CDKN1A expression remained unchanged (**Fig. 1F–H**). Concurrently, O. Tau increased expression of the inflammatory genes TNF, IL6, and IL1B (**Fig. 1I–K**). These effects were absent when heparin alone or the KRT8 control protein was used. Heparin reduced tau uptake, and markers linked to inflammation and senescence were lower when tau was combined with heparin than when tau was used alone.

### hTau mice show cell-cycle arrest and SASP-associated transcriptional changes in whole brain and astrocyte enriched fractions

To determine whether the transcriptional response observed in tau-treated astrocytes was also present in vivo, we measured cell-cycle regulatory and SASP-associated transcripts in 9-month-old hTau mice. In whole-brain tissue, expression of Cdkn2a, Cdkn1a, and Trp53 was higher in hTau mice than in WT littermates (**Fig. 2A–C**). The inflammatory transcripts Il1b and Il6 were also increased, whereas Ccl2 was not significantly altered (**Fig. 2D–F**). We next examined astrocyte-enriched fractions isolated from the same animals. Expression of Cdkn2a and Trp53 was higher in hTau fractions than in WT fractions (**Fig. 2H,I**). In contrast, Cdkn1a was lower in the hTau fractions (**Fig. 2G**), despite being increased in whole-brain tissue. This difference indicates that Cdkn1a is regulated differently in the astrocyte-enriched fraction and the broader brain tissue. Expression of the SASP-associated genes Il6, Tnf, and Serpine1 was also increased in hTau astrocyte-enriched fractions (**Fig. 2J–L**). Il1b was higher on average but did not reach statistical significance (P = 0.0552; **Fig. 2M**).

**Figure 2:**
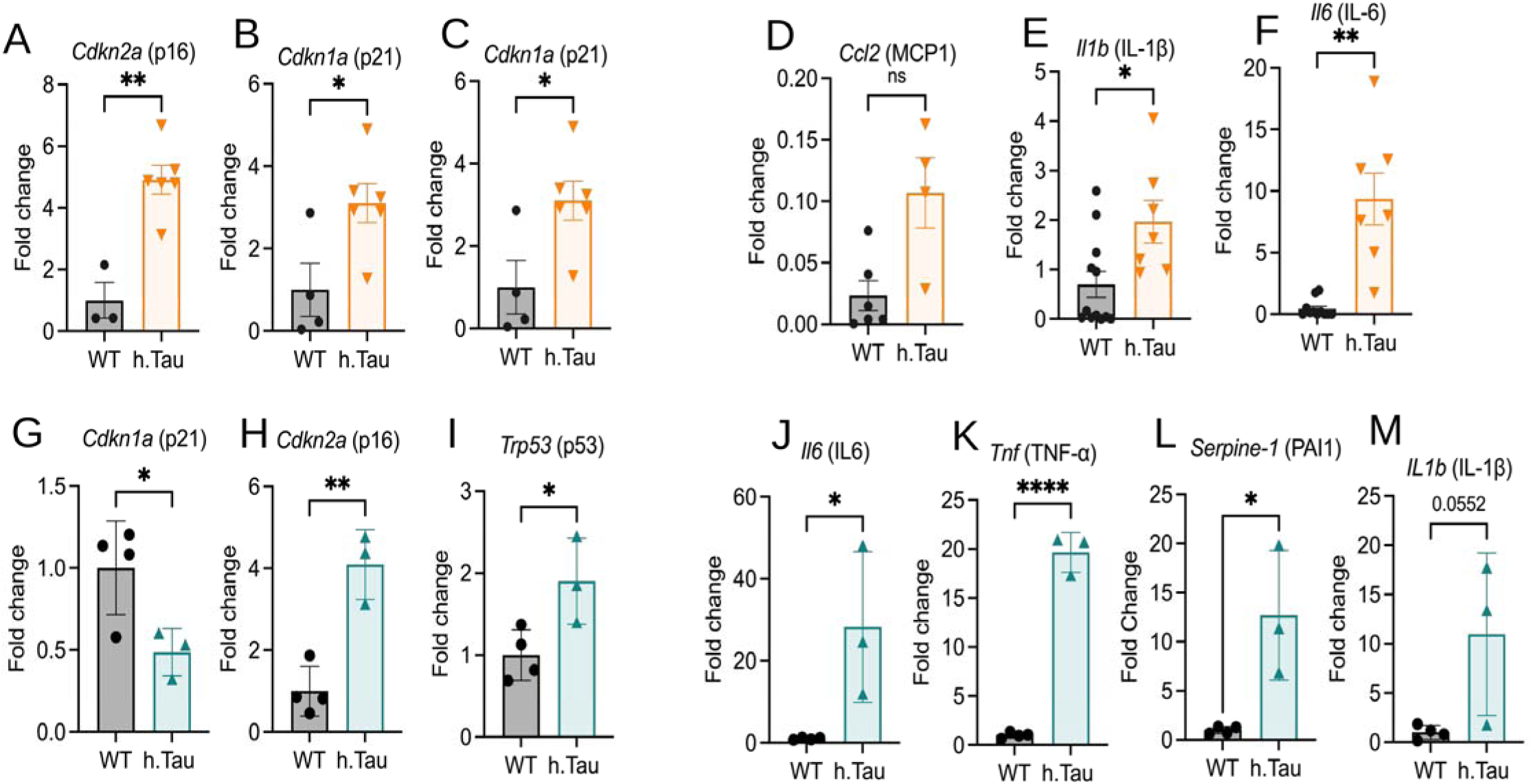
Cell-cycle-arrest and SASP markers are induced in hTau mouse whole brain and in astrocyte-enriched fractions. (**A-F**) qRT-PCR analysis of Cdkn2a/p16 (**A**), Cdkn1a/p21 (**B, C**), Ccl2/MCP-1 (**D**), Il1b/IL-10 (**E**), and Il6/IL-6 (**F**) in whole cortical/brain lysates from 9-month-old wild-type (WT) and hTau mice (n = 6-12**). (G-M**) qRT-PCR analysis of cell-cycle-arrest and SASP-associated markers in astrocyte-enriched fractions isolated from the same cohort by MACS using anti-ACSA-2 microbeads: Cdkn1a/p21 (**G**), Cdkn2a/p16 (**H**), Trp53/p53 (**I**), Il6/IL-6 (**J**), Tnf/TNF-a (**K**), Serpine1/PAI-1 (**L**), and II1b/IL-1p (**M**). SASP-associated transcripts and most cell-cycle-arrest markers were increased in hTau whole-brain samples and astrocyte-enriched fractions. Cdkn1a/p21 showed a different pattern, increasing in whole-brain lysate but trending lower in the astrocyte-enriched fraction. Data are mean ± SEM.; (n = 3-4) mice per group. Statistical comparisons were performed using two-tailed unpaired t-tests, with Welch’s correction applied when variances were unequal. *P < 0.05, **P < 0.01, ****P < 0.0001; ns, not significant. The exact P value is shown for panel M (P = 0.0552).

### OXPHOS and oxidative stress response signatures are reduced in hTau astrocytes

The gene set enrichment analysis showed that oxidative phosphorylation (OXPHOS) and the ROS/oxidative-stress response signatures were significantly negatively enriched in the hTau astrocytes as compared to the wild type (**Fig. 3A**). Oxidative phosphorylation showed the strongest negative enrichment (normalized enrichment score [NES] = –2.35, q < 0.001; **Fig. 3B**), followed by the ROS/oxidative-stress response (NES = –1.73, q = 0.004; **Fig. 3C**). Nevertheless, DNA repair, the inflammatory response, and cellular senescence did not reach statistical significance (q = 0.214, 0.122, and 0.496, respectively; **Fig. 3A**).

**Figure 3:**
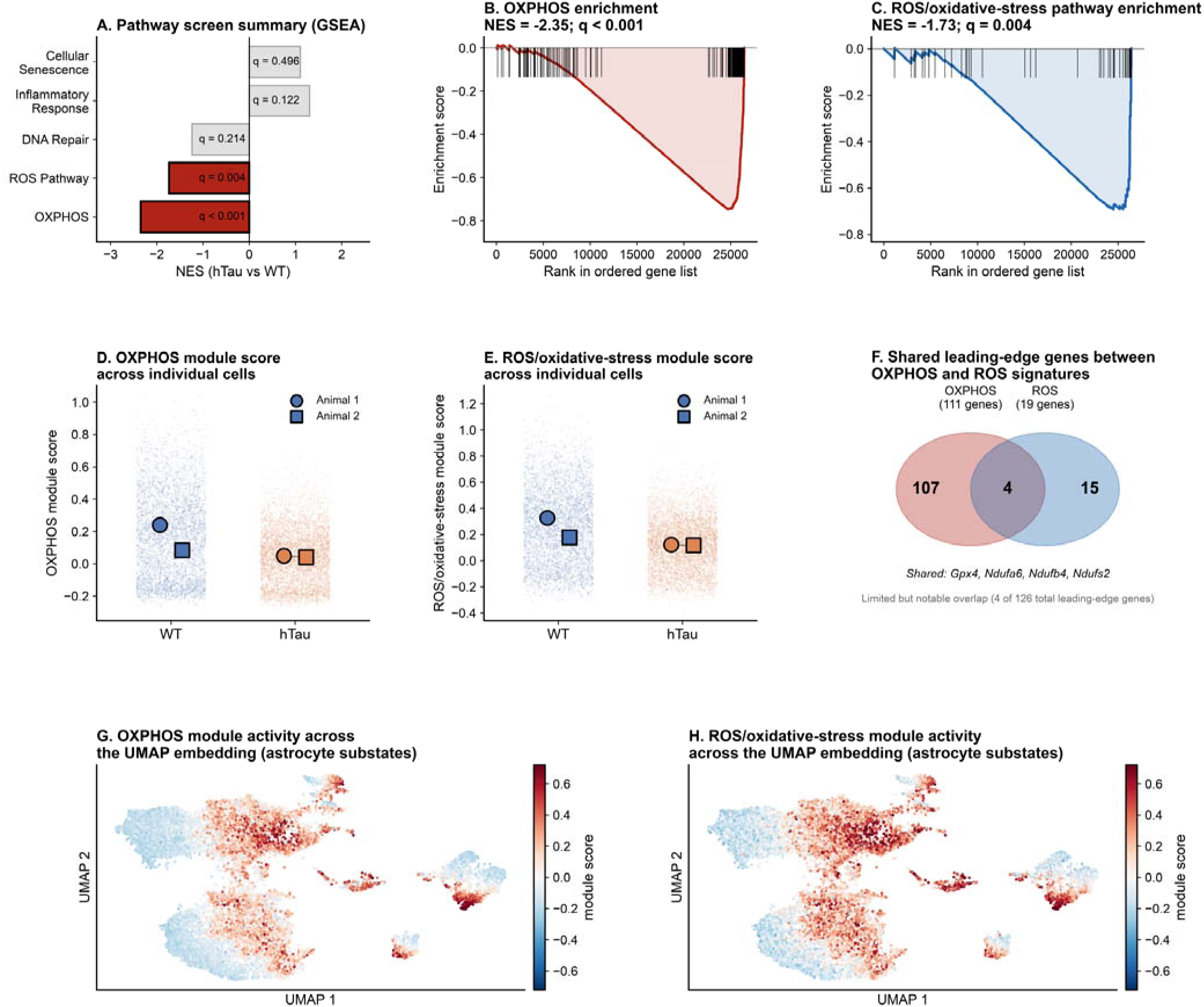
Oxidative phosphorylation and oxidative-stress response programs are suppressed in astrocytes from aged hTau mice. (**A**) Summary of the gene set enrichment analysis (GSEA) for five pathway categories comparing hTa**u** and WT astrocytes. The bars show normalized enrichment scores (NES); red indicates pathways meeting **t**he false discovery rate criterion of q < 0.05, and gray indicates pathways that are not significant. (**B and C**) Th**e r**unning enrichment scores for oxidative phosphorylation (OXPHOS; NES = –2.35, q < 0.001) (**B**) and reactive **ox**ygen species (ROS)/oxidative-stress response (NES = –1.73, q = 0.004) (**C**). Vertical ticks mark gene-set m**em**bers in the ranked gene list. (**D and E**) Per-cell module scores from the OXPHOS (**D**) and ROS/oxidative-s**tre**ss (**E**) leading-edge gene sets. Small points represent individual astrocytes; large circles and squares show mean scores for individual animals (n = 2 animals per genotype). Blue represents WT, and orange represen**ts** hTau. (**F**) Overlap between the OXPHOS and ROS/oxidative-stress leading-edge gene sets, consisting of 11**1** and 19 genes, respectively. Four genesGpx4, Ndufa6, Ndufb4, and Ndufs2are common to both sets, for a tot**al** of 126 unique genes. (G and H) OXPHOS (**G**) and ROS/oxidative-stress (**H**) module scores projected onto th**e** astrocyte UMAP embedding. Each point represents one cell and is coloured by its module score, usin**g** the same color scale in both panels. UMAP stands for uniform manifold approximation and projection.

To assess how these transcriptional changes are distributed, we computed module scores per cell using the leading-edge genes contributing to each enrichment signal. In the hTau astrocytes, both the OXPHOS and ROS/oxidative-stress scores were decreased, the mean scores being lower in each hTau animal than in either of the WT animals (**Fig. 3D and E**). Although the animal-level summaries agreed with the GSEA results, the comparison involved only two animals per genotype.

The OXPHOS set included 111 genes and the ROS/oxidative-stress set included 19 genes. Four genes (Gpx4, Ndufa6, Ndufb4, Ndufs2) were present in both sets, leaving 107 genes specific to OXPHOS and 15 unique to the ROS/oxidative-stress group (**Fig 3F**). The two enrichment signals were therefore derived from largely non-overlapping gene sets. Module scores plotted on the astrocyte UMAP embedding showed similar distributions, with overlapping regions of high and low scores across transcriptional substates (**Fig. 3G and 3H**).

### Soluble tau induces early mitochondrial oxidative and bioenergetic stress in primary human astrocytes

To assess whether mitochondrial changes precede cytoskeletal and senescence-associated responses to soluble tau, mitochondrial superoxide, ATP levels, and stress-response proteins were measured in primary human astrocytes over time. MitoTracker fluorescence decreased within 30 minutes of O. Tau exposure and gradually returned toward control levels over the next three hours (**Fig. 4A,B**). MitoSOX fluorescence increased at 30 minutes, indicating elevated mitochondrial superoxide (**Fig. 4C**). The signal remained slightly above control at several later time points but approached baseline by 24 hours. Treatment with antimycin A and rotenone caused a pronounced increase in MitoSOX fluorescence, confirming assay responsiveness (**Fig. 4C**). ATP levels also dropped within 30 minutes of O. Tau exposure and stayed below control levels for three hours (**Fig. 4D**). These changes occurred before detectable alterations in microtubule stability (**Fig. 5H**). This indicates that mitochondrial oxidative and bioenergetic stress is one of the earliest measured responses to soluble tau. Proteins involved in mitochondrial antioxidant defense and metabolic regulation were then examined. SOD2 abundance decreased four hours after O. Tau exposure, followed by reductions in HO-1 at eight hours and PGC-1a at 12 hours (**Fig. 4E-G**). The initial increase in mitochondrial superoxide and reduction in ATP therefore preceded decreases in these mitochondrial stress-response proteins. Mitochondrial changes also occurred before the induction of CDKN2A, CDKN1A, and TP53, which were not significantly altered until 24 hours (**Fig. 5E–G**). This temporal sequence places mitochondrial oxidative and bioenergetic stress before the detectable senescence-associated transcriptional response to soluble tau.

**Figure 4:**
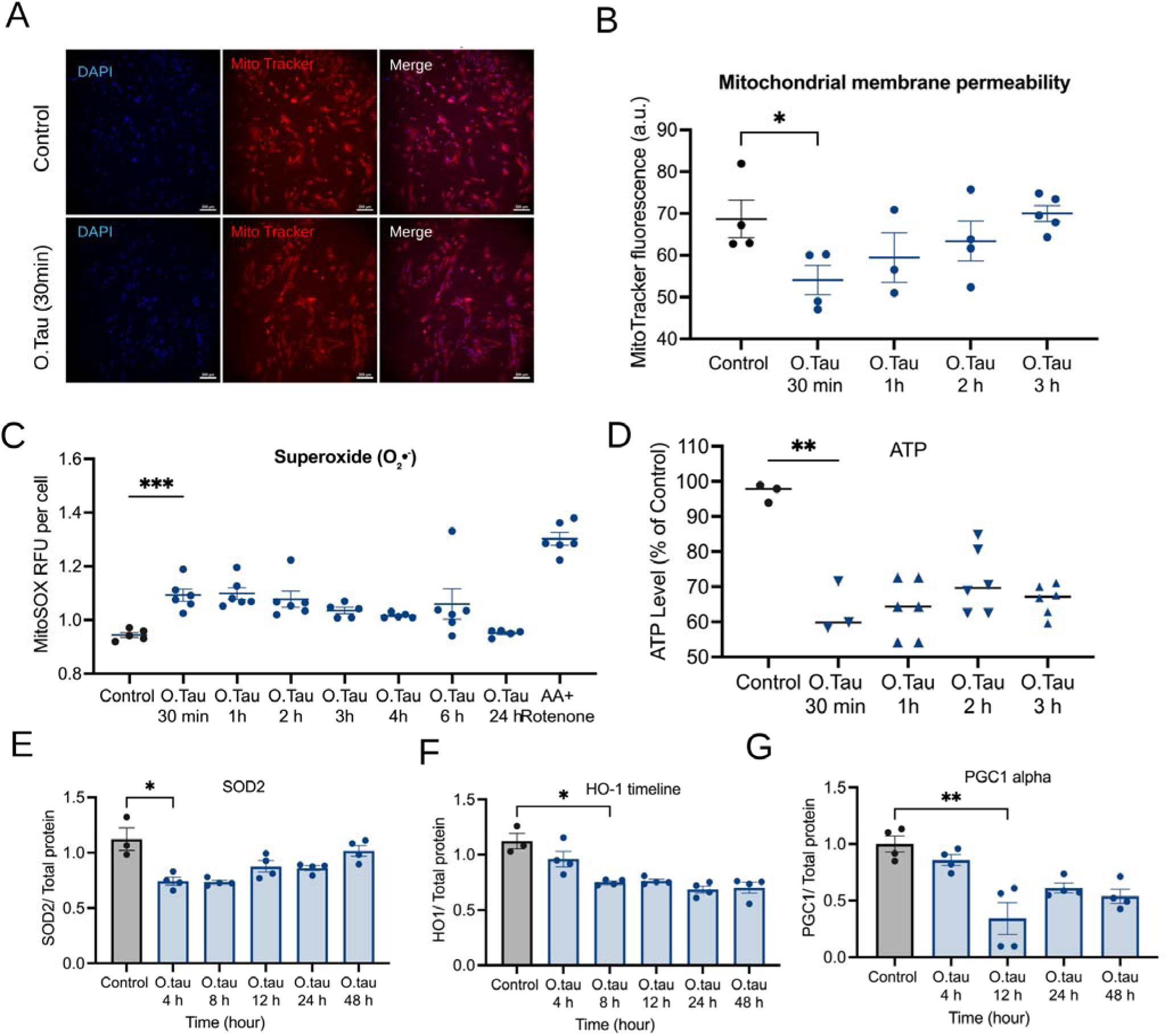
Soluble tau induces rapid mitochondrial dysfunction that precedes progressive remodeling of the mitochondrial stress-response and biogenesis program. (**A**) Representative epifluorescence images of astrocytes stained with DAPI (nuclei) and MitoTracker (mitochondria), along with merged images, for Control astrocytes and those exposed to O.Tau for 30 minutes. (**B**) Mitochondrial membrane integrity, assessed by MitoTracker fluorescence intensity, in Control astrocytes and those exposed to O.Tau for 30 minutes, 1 hour, 2 hours, or 3 hours (n = 4 independent experiments); *P < 0.05 at 30 minutes compared to Control, with values tending to return to baseline by 3 hours. (**C**) Mitochondrial superoxide (measured as MitoSOX Red fluorescence per cell) in Control and O.Tau-exposed astrocytes over 30 minutes to 24 hours, with antimycin A plus rotenone as positive control for maximal mitochondrial ROS (n = 2 independent experiment); ***P < 0.001 at 30 minutes versus Control (**D**) ATP levels (percentage of Control) during the same early time course (*P < 0.01 at 30 minutes**). (E-G**) Immunoblot quantification (normalized to total protein) of SOD2 (**E**), HO-1 (**F**), and PGC-1a (**G**) in astrocytes 4 to 48 hours after O.Tau exposure, all showing significant early reduction compared to Control (SOD2, P < 0.05 at 4 hours; HO-1, P < 0.05 at 8 hours; PGC-1a, *P < 0.01 at 12 hours). Data are mean ± SEM; n = 4 independent experiments at each time point. One-way ANOVA with correction for multiple comparisons versus Control (Methods). *P < 0.05, **P < 0.01, ***P < 0.001.

**Figure 5:**
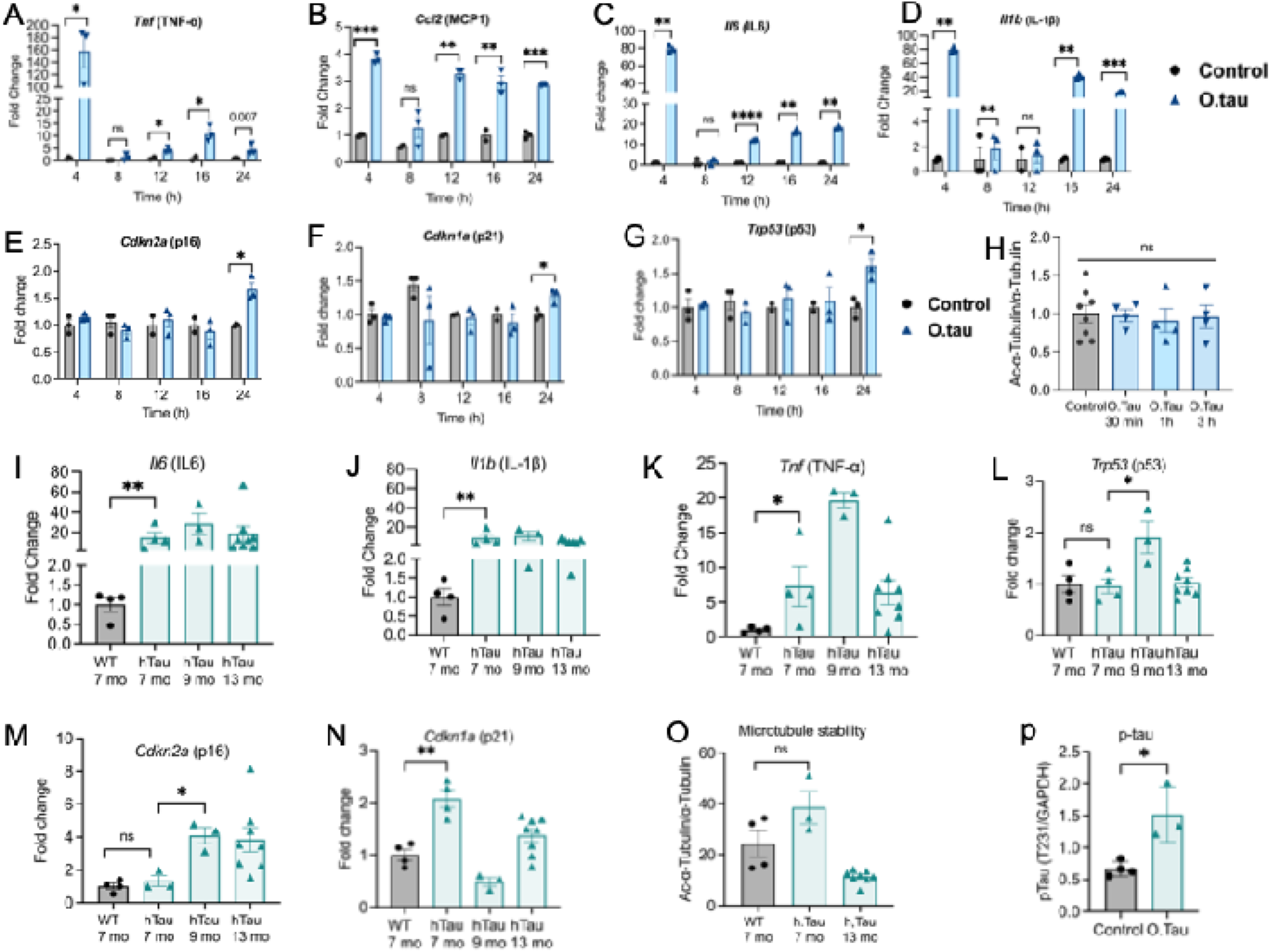
Tau elicits a biphasic astrocyte stress response early SASP cytokine induction followed by delayed senescence-marker expression that is recapitulated with age in hTau mice. (**A-D**), time course (4 to 24 hours) of SASP cytokine induction in primary human astrocytes exposed to O.Tau versus Control: Tnf/TNF-a (**A**), Ccl2/MCP-1 (**B**), II6/IL-6 (**C**), and Il1b/IL-10 (**D**). All four cytokines increased within 4 hours of tau exposure. E-G, over the same time course, cell-cycle-arrest markers Cdkn2a/p16 (**E**), Cdkn1a/p21 (**F**), and Trp53/p53 (**G**) did not differ from Control until 24 hours, showing SASP activation precedes senescence marker induction. (**H**), microtubule stability (acetylated-a-tubulin/total a-tubulin) in Control versus O.Tau-exposed astrocytes during 30 minutes to 3 hours was unchanged (ns), consistent with cytoskeletal destabilization requiring longer tau exposure than the acute bioenergetic and inflammatory responses in (**A-D). (I-P**), show the same temporal pattern is observed in vivo by age: Il6 (**I**), Il1b (**J**), and Tnf (**K**) were elevated in hTau compared to WT cortex/astrocyte-enriched tissue at 7 months; Trp53/p53 (**L**) and Cdkn2a/p16 (**M**) reached significance between 7 and 9 months; Cdkn1a/p21 (**N**) was elevated by 7 months. (**O**), microtubule stability (Ac-tubulin/tubulin) in WT (7 mo) and hTau (7 and 13 mo) cortex. (**P**), phosphorylated tau (pT231, normalized to GAPDH) was higher in hTau than control brain (P < 0.05), demonstrating tau pathology. Data are mean ± SEM. One-way ANOVA (in vitro) or two-tailed unpaired t-test / one-way ANOVA across ages (in vivo), with correction for multiple comparisons. *P < 0.05, **P < 0.01, ***P < 0.001, ****P < 0.0001; ns, not significant; exact value shown for A at 24 hours (P = 0.007).

### SASP cytokine induction precedes senescence-marker expression in a biphasic astrocyte response that recurs with age in hTau mice

Since soluble tau rapidly disrupted mitochondrial integrity, increased mitochondrial superoxide, and decreased ATP levels within 30 minutes (**Fig. 4A-D**), we investigated how these early mitochondrial changes relate to inflammatory activation, cell-cycle arrest, and microtubule stability. Primary human astrocytes were exposed to O.Tau over 4 to 24 hours, and levels of SASP-associated cytokines, senescence markers, and microtubule stability were measured during this period.

Exposure of primary Human astrocytes to O. Tau triggered a rapid inflammatory response. TNF was raised as early as 4 hours and stayed high at later time points, with significant increases observed at 4, 12, and 16 hours and a continuing rise at 24 hours (**Fig. 5A**). Ccl2/MCP1 also increased by 4 hours, showed no significant change at 8 hours, and was once more elevated from 12 to 24 hours (**Fig. 5B**). Il6 exhibited a marked early induction at 4 hours, followed by significant increases from 12 to 24 hours (**Fig. 5C**). Il1b was elevated at both 4 and 8 hours, was not significantly different at 12 hours, and then increased again at 16 and 24 hours (**Fig. 5D**). Hence, exposure to tau led to the activation of SASP-associated inflammatory genes before the overt activation of cell-cycle arrest markers.

Markers of cell-cycle arrest were delayed. Cdkn2a/p16, Cdkn1a/p21, and Trp53/p53 did not differ from control at early time points but increased by 24 hours after O.Tau exposure (**Fig. 5E-G**). Microtubule stability, measured by acetylated to total alpha-tubulin ratio, showed no significant change between 30 minutes and 3 hours (**Fig. 5H**).

We then investigated whether this temporal pattern was reproduced as tau pathology advanced in the hTau mice. At 7 months of age the SASP-associated transcripts were already upregulated in the hTau mice, involving Il6, Il1b, and Tnf (**Fig. 5I-K**). In contrast, the cell-cycle arrest markers displayed a more delayed or marker-specific pattern: Trp53/p53 was not significantly altered at 7 months but did increase at 9 months (**Fig. 5L**), Cdkn2a/p16 was not significantly changed at 7 months but rose at 9 months (**Fig. 5M**), and Cdkn1a/p21 was increased at 7 months (**Fig. 5N**). Microtubule stability was also examined across the age groups (**Fig. 5O**). The increased level of phosphorylated tau at T231 demonstrated the presence of tau-associated pathology in the respective samples (**Fig. 5P**).

When the data are considered as a whole, they indicate that astrocytes respond to tau in a stepwise manner: mitochondrial oxidative and bioenergetic stress occurs first (**Fig. 4A-D**), prior to any detectable microtubule destabilization (**Figure 5H**) and prior to the activation of the senescence-associated cell-cycle arrest program. This is then followed by the early induction of cytokines associated with the SASP (**Figures 5A-D**), while the markers for microtubule destabilization and cell-cycle arrest appear later (**Figures 5E-H**).

### Soluble tau impairs mitochondrial respiration in primary human astrocytes

Since O.Tau caused a rapid reduction in mitochondrial membrane integrity, an increase in mitochondrial superoxide, and a depletion of ATP (**Fig. 4A-D**), we tested whether these early mitochondrial abnormalities led to a detectable defect in oxidative metabolism. We measured oxygen consumption rate during a mitochondrial stress test in primary human astrocytes treated with O.Tau.

Control astrocytes showed the expected OCR profile, with measurable basal respiration, decreased OCR after ATP synthase inhibition, and increased respiration after mitochondrial uncoupling (**Fig. 6A**). In contrast, astrocytes exposed to O.Tau had lower OCR throughout the assay, indicating reduced mitochondrial respiratory activity (**Fig. 6A**). Quantification showed O.Tau significantly decreased basal respiration (**Fig. 6B**), indicating reduced mitochondrial oxygen consumption at rest. ATP-linked respiration was also decreased after O.Tau exposure (**Fig. 6C**), consistent with impaired mitochondrial ATP production. O.Tau also decreased proton leak (**Fig. 6D**), indicating a general suppression of mitochondrial oxygen flux beyond ATP-coupled respiration. Maximal respiration was significantly reduced in O.Tau-exposed astrocytes (**Fig. 6E**), showing impaired maximal respiratory capacity under uncoupled conditions. Thus, tau exposure hampers both basal mitochondrial respiration and astrocytes’ ability to increase oxidative metabolism in response to energy demand.

**Figure 6:**
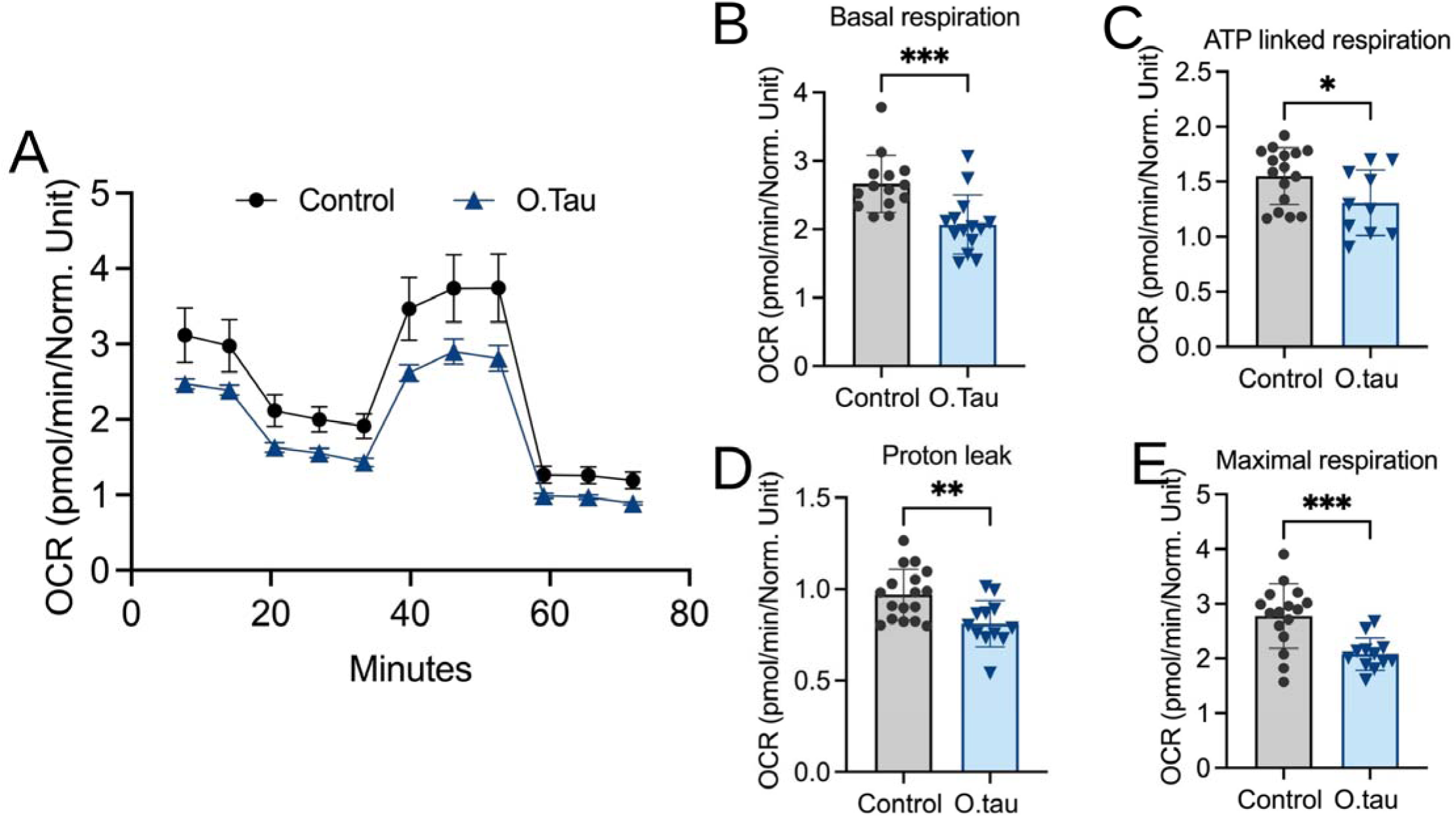
Soluble tau impairs mitochondrial oxidative respiration in astrocytes. (**A**) Oxygen consumption rate (OCR) traces from a mitochondrial stress test (Seahorse extracellular flux analysis) in control cells versus those exposed to O.Tau among primary human astrocytes, following sequential injections of oligomycin, FCCP, and rotenone/antimycin A. (**B to E):** Quantification of basal respiration (B, ***P < 0.001), ATP-linked respiration (**C, P < 0.05**), proton leak (**D, *P < 0.01**), and maximal respiration (**E, ***P < 0.001**), all reduced in O.Tau-exposed astrocytes compared to controls. Data are mean ± SD.; n = 4 independent wells/replicates per group, from n = 3-4 independent experiments. A two-tailed unpaired t-test was used. *P < 0.05, **P < 0.01, ***P < 0.001.

Combined with early defects in mitochondrial membrane, ROS, and ATP (**Fig. 4**), these findings show soluble tau rapidly disrupts astrocyte mitochondrial function and causes prolonged impairment of oxidative respiratory capacity.

### Mitochondrial ROS scavenging with MitoTEMPO suppresses tau-induced astrocyte senescence and SASP activation in vivo

Since soluble tau caused a rapid increase in mitochondrial superoxide and reduced mitochondrial function in astrocytes, we next investigated whether mitochondrial ROS contributes to the senescence– and SASP-associated astrocyte response in vivo. The hTau mice received daily intraperitoneal injections of the mitochondria-targeted antioxidant MitoTEMPO at 1.5 mg/kg for two weeks. brains were then removed, dissociated, and the astrocytes were enriched using MACS sorting with anti-ACSA-2 microbeads for subsequent transcript analysis (**Fig. 7A**). In astrocyte-enriched fractions, hTau mice showed higher Cdkn2a/p16 expression than WT (**Fig. 7B**). MitoTEMPO treatment significantly decreased Cdkn2a/p16 in hTau mice to WT levels (**Fig. 7B**). There was no significant difference in Trp53/p53 or Cdkn1a/p21 among groups, and MitoTEMPO had no effect on these markers (**Fig. 7C,D**). These data show mitochondrial ROS scavenging specifically reduces the p16 component of the cell-cycle arrest response in astrocytes. We examined SASP-associated transcripts in the same astrocyte-enriched fractions. Il6 was higher in hTau mice than WT and was significantly decreased by MitoTEMPO treatment (**Fig. 7E**). Tnf was also elevated in hTau mice, but MitoTEMPO did not significantly reduce Tnf expression (**Fig. 7F**). Serpine1/PAI-1 showed no significant changes among groups (**Fig. 7G**).

**Figure 7:**
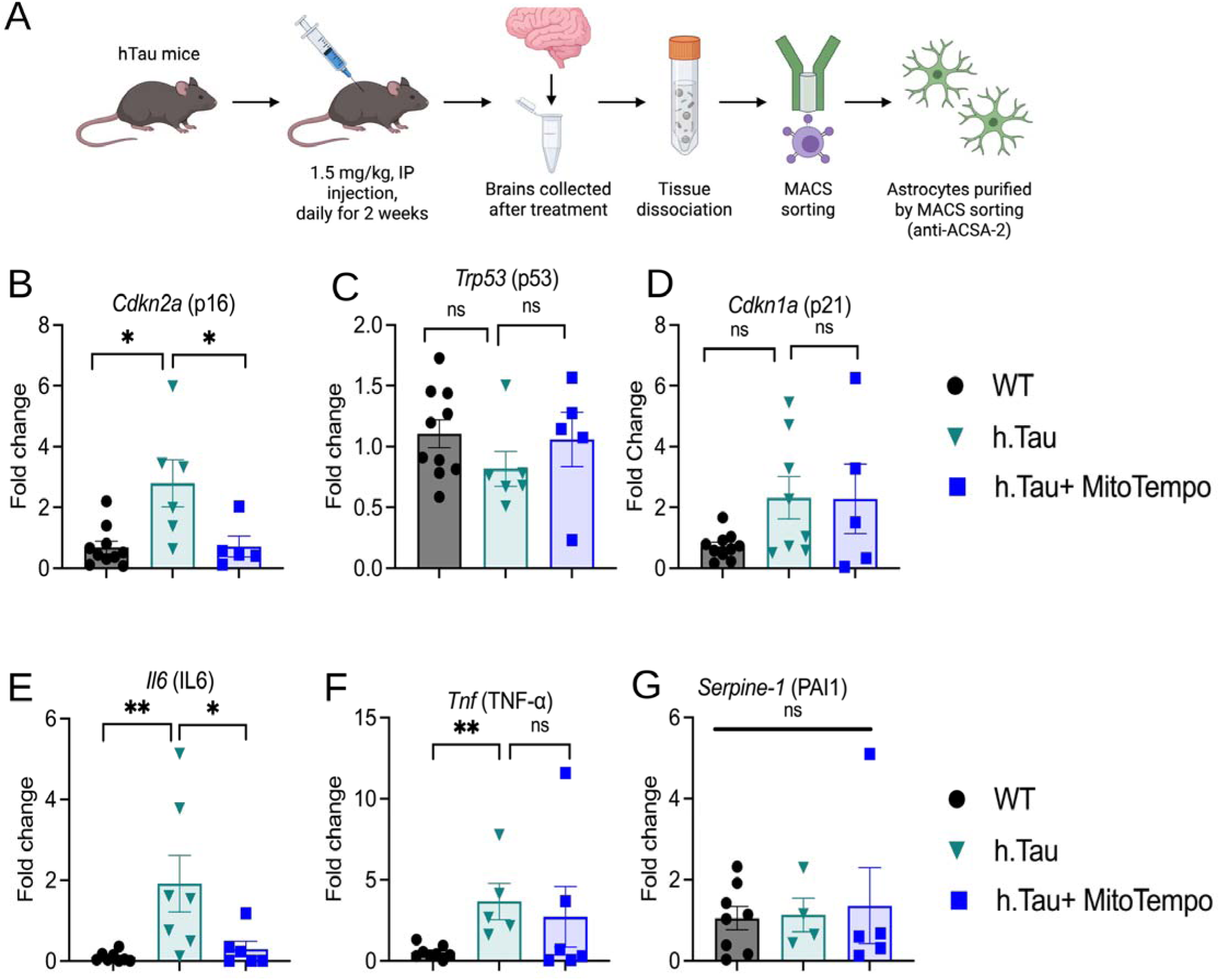
Astrocyte-targeted mitochondrial ROS scavenging with MitoTEMPO suppresses tau-induced senescence and SASP activation in vivo. (**A**) Experimental schematic the hTau mice were given either MitoTEMPO or the vehicle (1.5 mg kg^−1^, by intraperitoneal injection, once a day for 2 weeks); after that, the brains were collected, dissociated, and the astrocytes purified using magnetic-activated cell sorting (MACS) with anti-ACSA-2 microbeads for further analysis. **B-D:** The fold change (by qRT-PCR) of the cell-cycle-arrest markers Cdkn2a/p16 (**B**), Trp53/p53 (**C**), and Cdkn1a/p21 (**D**) in the astrocytes from WT, hTau, and hTau + MitoTEMPO mice. MitoTEMPO significantly reduced the hTau-induced increase in Cdkn2a/p16 (P < 0.05) but had no effect on Trp53 or Cdkn1a (both not significant). (**E-G**) The fold change of the SASP markers II6/IL-6 (**E**), Tnf/TNF-a (**F**), and Serpine1/PAI-1 (**G**). MitoTEMPO significantly reduced the hTau-associated increase in Il6 (P < 0.05) but did not significantly reduce Tnf or Serpine1 (not significant). The data are presented as mean ± SEM.; the number of animals per group is n = 5-10. A one-way ANOVA was carried out with correction for multiple comparisons among the WT, hTau, and hTau + MitoTEMPO groups (Methods). *P < 0.05, **P < 0.01; ns, not significant

Together, the data indicate that scavenging mitochondrial ROS in hTau mice reduces certain astrocyte senescence/SASP responses, notably Cdkn2a/p16 and Il6. The lack of a significant decrease in Trp53, Cdkn1a, Tnf, and Serpine1 indicates that MitoTEMPO does not globally suppress all senescence or SASP transcripts but affects only a subset of the tau-related astrocyte response.

### Tau-induced senescent astrocytes impair neuronal dendritic architecture through secreted factors

Astrocytes support synaptic structure and function^56^, whereas neurotoxic reactive astrocyte states can damage neurons^57^. Since exposure of astrocytes to O.Tau led to senescence and the elevation of SASP-associated markers, we investigated whether astrocytes exposed to tau were able to impair the structure of neurons in a non-cell-autonomous way. At 18 days in vitro, primary astrocytes were exposed to soluble tau aggregates or vehicle, then co-cultured for 96 hours with primary mouse cortical neurons prior to morphological analysis (**Fig. 8A**).

**Figure 8:**
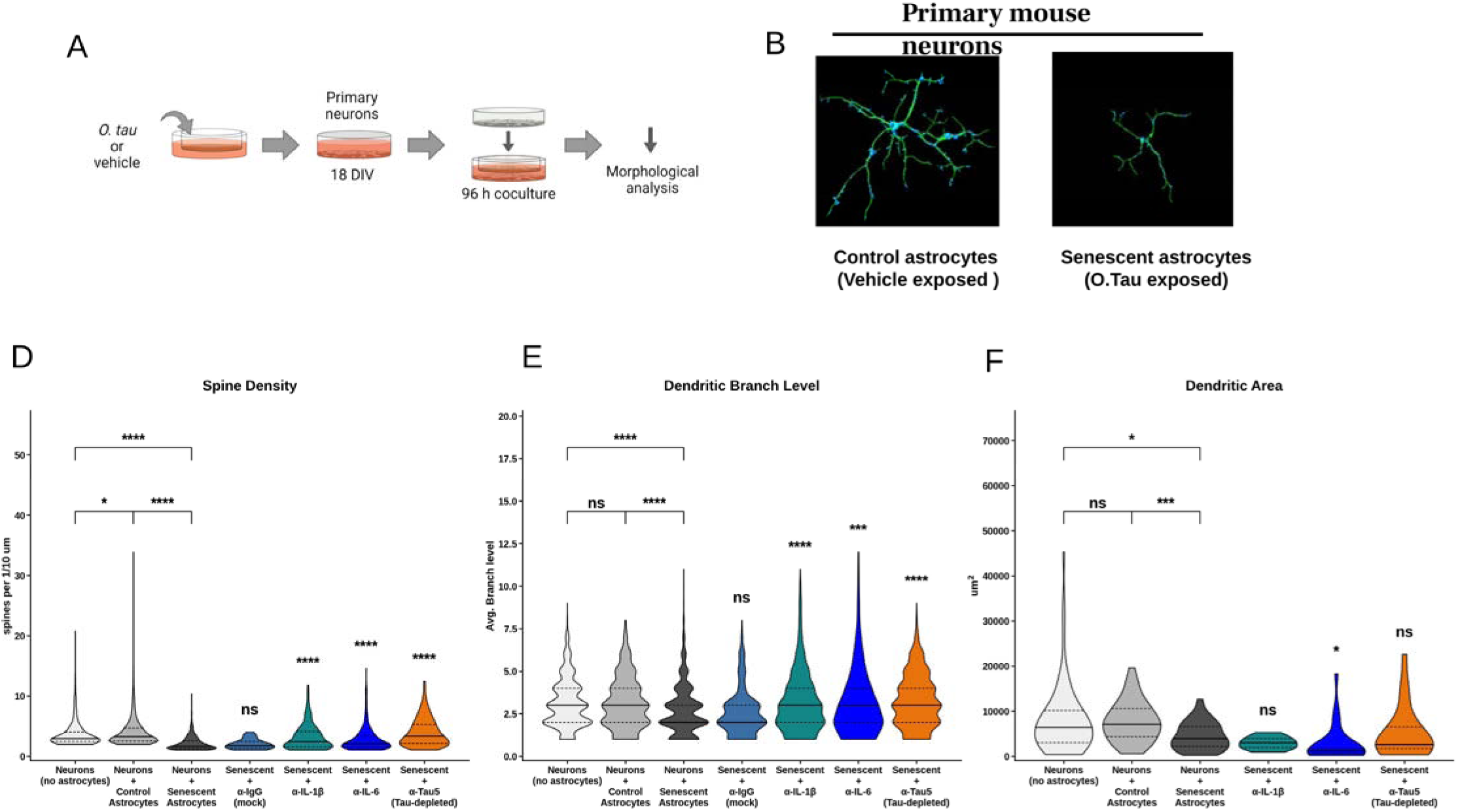
Tau-induced senescent astrocytes impair neuronal dendritic architecture through a tau-dependent secreted factor. (**A**) Schematic showing the coculture approach: astrocytes that had been exposed to O.Tau or had received vehicle alone were combined with primary mouse cortical neurons (which had been grown in vitro for 18 days) and then left to coculture for 96 hours before undergoing morphological analysis. (**B**), Example images of the neurons that were co-cultured with control (vehicle-exposed) astrocytes as compared to those co-cultured with senescent (O.Tau-exposed) astrocytes, showing a decrease in dendritic arborization when the neurons were co-cultured with senescent astrocytes. (**D-F**), Violin plots (showing the median and quartiles) of dendritic spine density (**D**; number of spines per 10 µm of dendrite), average dendritic branch level (**E**), and dendritic area (F; in µm²) for seven different conditions: neurons grown alone (Neurons, no astrocytes; n = 429, 2,106 and 80 dendrite segments per neuron for spine density, branch level and dendritic area, respectively), neurons cocultured with vehicle-treated control astrocytes (Neurons + Astrocytes; n = 2,843, 1,130 and 74), neurons cocultured with tau-exposed, senescent astrocytes (Neurons + Senescent Astrocytes; n = 946, 3,498 and 92), and neurons cocultured with senescent-astrocyte-conditioned media from which IgG (mock), IL-1p, IL-6 or tau (a-Tau5) had been immunodepleted before application (n = 17 to 25 cells per arm for dendritic area; Methods). Black brackets indicate the comparisons between Neurons, Control Astrocytes, and Senescent Astrocytes; asterisks above each depletion condition show the comparison with Neurons + Senescent Astrocytes within the same test group. The Kruskal-Wallis test with Dunn’s multiple-comparisons test (Bonferroni-corrected). *P < 0.05, ***P < 0.001, ****P < 0.0001; ns, not significant.

Neurons co-cultured with vehicle-exposed astrocytes maintained complex dendritic arbors, but those with tau-exposed astrocytes showed visibly decreased dendritic complexity (**Fig. 8B**). Quantification showed co-culture with senescent astrocytes reduced dendritic spine density, branch level, and area compared to neurons grown alone or with vehicle-treated astrocytes (**Fig. 8D-F**). These results show astrocytes exposed to tau impair multiple aspects of neuronal dendritic architecture, including spine density, arbor complexity, and dendritic field size.

To identify the secreted factors responsible for this effect, we removed IL-1p, IL-6, or tau from the conditioned media of senescent astrocytes using antibodies before applying the conditioned media to neurons, along with a control experiment in which IgG was used for depletion. The IgG (mock depletion) had no effect on spine density, branch level, or dendritic area, indicating that neuronal impairment persisted even after the control immunodepleting (**Fig. 8D-F**). Only the depletion of tau, not that of IL-1p or IL-6, led to a rescue of dendritic spine density (**Fig. 8D**). Dendritic branch level was partially restored by the depletion of tau, IL-1p, or IL-6, whereas mock depletion did not result in any improvement in this parameter (**Fig. 8E**). None of the depletion conditions had a significant effect on dendritic area, leaving the basis of the residual deficit unresolved (**Fig. 8F**).

Taken as a whole, the data show that astrocytes containing tau impair the structural integrity of neurons by means of soluble factors in the astrocyte secretome. It seems that tau in the secretome is the main factor leading to a loss of dendritic spines, whereas both tau, IL-1p, and IL-6 contribute to a decrease in dendritic branch complexity. The fact that there is no rescue of the dendritic area indicates that the more general structural deficit cannot be completely accounted for by any of the factors tested.

### Alterations in astrocyte vascular signaling and neurovascular responses in hTau mice

An analysis of intercellular signaling showed extensive changes in the inferred communication between astrocytes and various vascular cell populations in the hTau mice (**Fig. 9A-C**). Among the source-target cell populations examined, the number of significant interactions rose, including bidirectional interactions between astrocytes and endothelial cells, pericytes, and vascular leptomeningeal cells. This suggests more widespread inferred signaling at the astrocyte-vascular interface, although the increase in interaction counts does not prove that communication is stronger or more effective.

**Figure 9.** Alterations in astrocyte vascular signaling and neurovascular responses in hTau mice. (**A**) A schematic illustration of the interactions between astrocytes and other cell types in the neurovascular unit: neuronal, endothelial, vascular smooth muscle/pericyte, and microglial cells. (**B**) A heatmap showing changes in significant intercellular signaling interactions compared with WT. Rows show source cell populations and columns show target populations. Values indicate the number of interactions differing with a permutation P-value less than 0.01**. (C**) The number of significant interactions in WT and hTau groups for specific directional astrocyte-vascular cell pairs. (**D**) Examples of signaling pairs showing increased or decreased signaling in hTau. Bars show how many directional astrocyte-vascular cell-pair contexts exhibit each change type (permutation P < 0.01, uncorrected). (**E and F**) Astrocyte-vascular signaling pairs related to neurovascular coupling (**E**) and disease (**F**) across specified source target cell populations. Dot size corresponds to permutation P value; color intensity reflects interaction strength based on mean expression**. (G**) Net directional cell-pair scores for signaling pairs grouped by neurovascular regulatory processes. Positive and negative scores indicate predominant increase and decrease in signaling, respectively. Groups were determined using literature-based annotations and do not constitute formal database enrichment.

Although there was an overall rise in the number of interactions, individual signaling pairs showed opposite trends (**Fig. 9D**). PTPN11-GSK3B increased in five directional astrocyte-vascular cell-pair situations, while HGF-MAP2K1, KMT2E-RARA, LY96-IKBKB, FZD8-LRP6, and EGF-CDK12 each increased in four contexts. Conversely, BCAN-NRCAM and SORL1-CNTFR decreased in four contexts, and VEGFA-GRIN2B, NRG1-ITGAV, TTR-MAPK3, and NTN3-DCC decreased in three contexts (permutation P < 0.01, uncorrected). These figures show the distribution of each type of change across different cell-pair contexts rather than the extent of the change, indicating that the general increase in inferred interactions occurred alongside specific losses in signaling.

The neurovascular coupling-related and disease-associated signaling pairs also varied in their distribution across the source target populations (**Fig. 9E and F**). While some pairs appeared in multiple directional contexts, others exhibited more limited patterns. These diagrams show both the strength of the interactions and the results of the permutation tests, providing clearer context for the cellular distribution of the selected signals.

When the signaling pairs were grouped based on literature, growth-factor signaling had the highest positive net directional score, while the scores for chemokine/inflammatory signaling and Wnt signaling related to vascular stability were smaller but positive (**Fig. 9G**). Extracellular matrix/adhesion had the most negative score, followed by cell guidance/trophic signaling and calcium-related processes. These descriptive scores indicate the balance between increased and decreased cell-pair contexts in each group; they do not represent formal pathway enrichment. Taken together, the results show a reorganization of inferred communication between astrocytes and blood vessels in the hTau mice, with increased growth-factor and inflammation-associated signaling and selective reductions in signaling related to extracellular organization and cell guidance.

### Astrocyte-targeted SOD2 overexpression shifts the profile of tau-induced deficits in evoked cerebral blood flow

Tau exposure induced mitochondrial oxidative stress in astrocytes and mitochondrial ROS scavenging attenuated selected senescence/SASP markers in vivo. We next asked whether increasing mitochondrial antioxidant capacity in astrocytes specifically could improve neurovascular function in hTau mice. We overexpressed the mitochondrial matrix antioxidant enzyme SOD2 using an astrocyte-targeted AAV vector^58,59^, with GFP-expressing hTau littermates as the controls. Whisker-stimulation-evoked cerebral blood flow (CBF) responses were measured by laser Doppler flowmetry under sequential superfusion with artificial cerebrospinal fluid (aCSF), the neuronal nitric oxide synthase inhibitor (NPA), and the pan-nitric oxide synthase inhibitor (L-NAME) (**Fig. 10A-C**).

**Figure 10:**
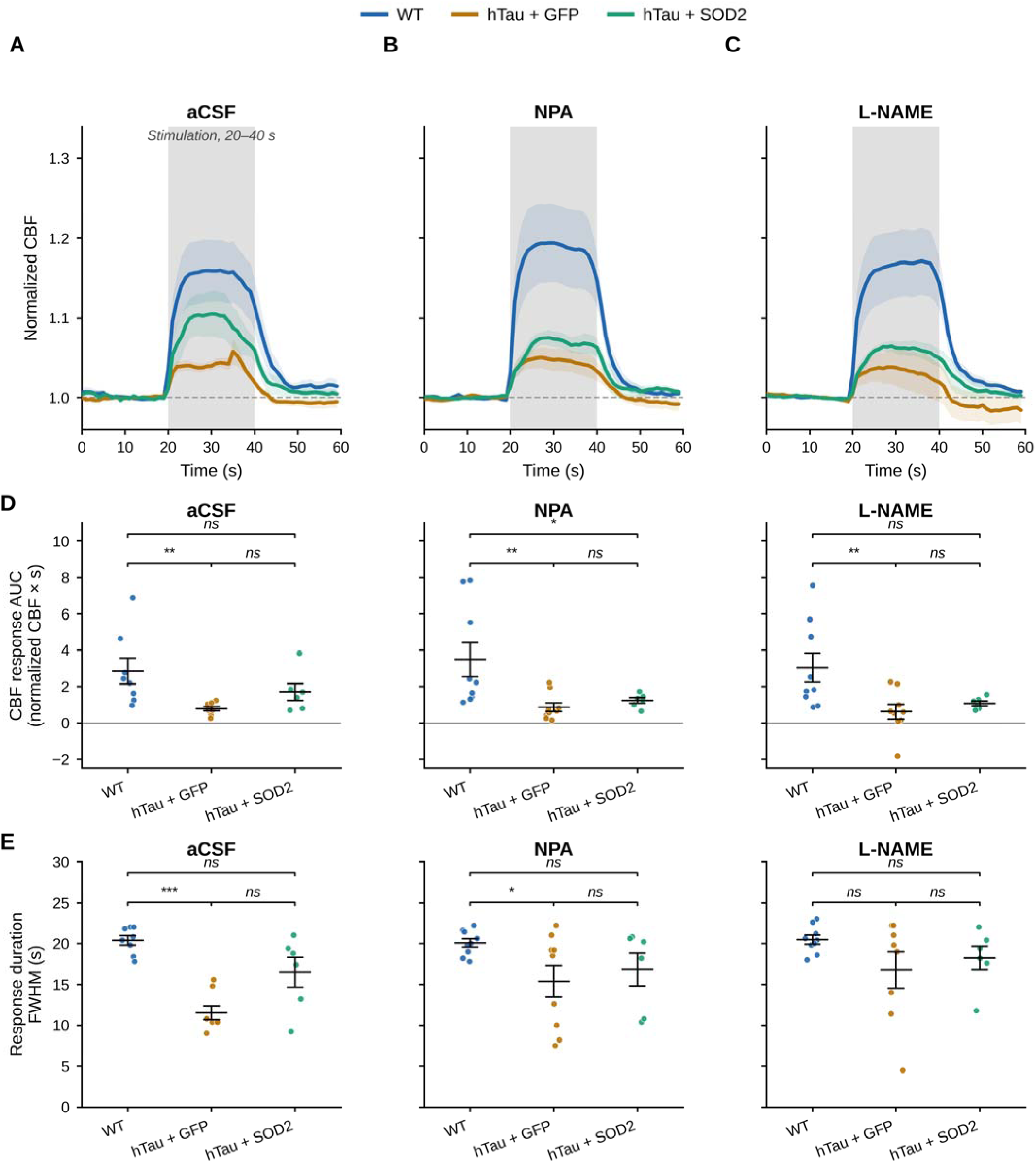
Astrocyte-targeted SOD2 overexpression partially normalizes tau-induced deficits in evoked cerebral blood flow. (**A-C**), Normalized cerebral blood flow (CBF) traces (mean ± SEM., shaded band) in WT, hTau + GFP, and hTau + SOD2 mice in response to whisker stimulation (20-40 s, gray shading), under sequential super fusion with artificial cerebrospinal fluid (aCSF; A), the neuronal nitric oxide synthase inhibitor NPA (**B**), and the pan-nitric-oxide-synthase inhibitor L-NAME (**C). (D**), CBF response area under the curve (AUC; normalized CBF × s) for each condition. hTau + GFP mice showed significantly reduced AUC relative to WT under all three conditions (aCSF, **P < 0.01; NPA, **P < 0.01; L-NAME, **P < 0.01); hTau + SOD2 mice did not differ significantly from WT under aCSF or L-NAME (ns) but remained significantly reduced under NPA (*P < 0.05) and did not differ significantly from hTau + GFP under any condition (ns). (**E**), Response duration (full width at half maximum, FWHM). hTau + GFP mice showed significantly shorter response duration than WT under aCSF (***P < 0.001) and NPA (*P < 0.05) but not L-NAME (ns); hTau + SOD2 mice did not differ significantly from WT or from hTau + GFP under any condition (ns). Data are mean ± SEM.; n = 17 mice/group. Repeated-measures mixed-effects model with multiplicity-adjusted P values for pairwise comparisons within each condition. *P < 0.05, **P < 0.01, ***P < 0.001; ns, not significant.

Across all three superfusion conditions, hTau + GFP mice showed reduced CBF response area under the curve (AUC) relative to WT mice (aCSF, NPA, and L-NAME: all P < 0.01; **Fig. 10D**), this demonstrates impaired evoked neurovascular responses in hTau mice. hTau + GFP mice also showed shorter response duration under aCSF (P < 0.001) and NPA (P < 0.05), but not under L-NAME (**Fig. 10E**). These data indicate that hTau mice have NVC deficits in both response magnitude and response duration, with the duration deficit most evident before complete nitric oxide synthase blockade.

Astrocyte-targeted SOD2 overexpression shifted CBF response magnitude toward the WT levels. hTau + SOD2 mice no longer differed significantly from WT under aCSF or L-NAME, although they remained significantly reduced relative to WT under NPA (P < 0.05; **Fig. 10D**). Response duration in hTau + SOD2 mice did not differ significantly from WT under any superfusion condition (**Fig. 10E**). However, hTau + SOD2 mice also did not differ significantly from hTau + GFP mice for either AUC or response duration under any condition. Therefore, the SOD2 effect should be interpreted as partial normalization of the response profile, or a shift toward WT values, rather than a statistically significant rescue. In conjunction with the MitoTEMPO data, these results motivate further investigation of astrocyte mitochondrial oxidative stress in neurovascular dysfunction. Nonetheless, astrocyte-specific SOD2 overexpression did not produce a statistically significant improvement relative to the hTau + GFP group in this experimental context.

## Discussion

Astrocytes play a critical role in supporting neurons and the cerebral vasculature; however, the mechanisms by which direct exposure to pathogenic tau leads to astrocyte dysfunction in tauopathy are not fully understood. The present study demonstrates that soluble tau aggregates are transmitted to primary human astrocytes via a heparin-sensitive mechanism, consistent with the previously described heparan sulfate proteoglycan (HSPG)-dependent uptake of tau in neurons^14^. Alternative entry pathways, such as low-density lipoprotein receptor-related protein 1 (LRP1), may also contribute^15^ (**Fig. 1A**). Exposure to tau rapidly increases mitochondrial oxidative stress, reduces ATP levels, and lowers the abundance of SOD2, HO-1, and PGC-1a (**Fig. 4**). These alterations occur prior to any detectable microtubule destabilization or induction of the cell-cycle-arrest markers assessed (**Fig. 5**). At subsequent time points, tau further decreases basal respiration, ATP-linked respiration, and maximal respiration (**Fig. 6**). Collectively, these results indicate that mitochondrial oxidative and bioenergetic stress represent early features of the astrocyte response to soluble pathogenic tau transmission, followed by a reduction in mitochondrial respiratory capacity. This sequence is consistent with evidence that mitochondrial oxidative stress contributes to the initiation and maintenance of cellular senescence and the senescence-associated secretory phenotype (SASP)^51–53,60,61^.

Cellular senescence is generally characterized by stable cell-cycle arrest and the development of a senescence-associated secretory phenotype (SASP)^38–44^. In the present study, inflammatory gene transcription occurred prior to any detectable induction of p16 in astrocytes exposed to tau. Within four hours of tau exposure, transcript levels of Tnf, Il6, Ccl2/MCP-1, and Il1b increased, whereas Cdkn2a/p16 did not (**Fig. 5A-E**). A comparable pattern was observed in aged hTau mice: Il6, Il1b, and Tnf were elevated by seven months, while Cdkn2a/p16 did not reach significance until nine months (**Fig. 5I-M**). Furthermore, SASP-associated transcript levels were higher in astrocyte-enriched fractions than in whole brain samples (**Fig. 2**). These findings suggest that p16 alone may not sufficiently capture the early inflammatory response of astrocytes to tau. As these experiments assessed transcript abundance, they do not establish the onset of stable cell-cycle arrest or determine whether the early transcriptional response is accompanied by cytokine secretion.

We examined the role of mitochondrial oxidative stress in mediating inflammatory and senescence-related responses in vivo. MitoTEMPO significantly reduced the increases in Il6 and Cdkn2a/p16 associated with hTau in astrocyte-enriched fractions but did not significantly affect Trp53/p53, Cdkn1a/p21, Tnf, or Serpine1 (**Fig. 7**). This selective effect indicates that mitochondrial oxidative stress contributes to specific aspects of the response rather than broadly suppressing transcripts associated with senescence and the SASP. These findings align with previous studies showing mitochondrial antioxidants protect in models of Alzheimer’s disease and vascular dysfunction^62–64^. Consistent with mitochondrial alterations observed in primary astrocytes exposed to tau, single-cell RNA sequencing revealed reduced expression of genes involved in mitochondrial electron transport, oxidative phosphorylation, and mitophagy in astrocytes from aged hTau mice (**Fig. 3**). Although these transcriptional changes do not directly measure mitochondrial function, their alignment with in vitro findings supports the hypothesis that mitochondrial pathways are disrupted in hTau astrocytes.

Our previous work demonstrated that soluble pathogenic tau enters brain microvascular endothelial cells, inducing cellular senescence and microvascular dysfunction^54^. The current findings extend the effects of soluble tau to astrocytes, identifying an additional cell type within the neurovascular unit that responds to tau. In isolated primary astrocytes, extracellular tau alone was sufficient to induce mitochondrial changes that led to inflammatory transcription, even without neurons or microglia. Thus, astrocytes can respond directly to extracellular tau, although the extent to which this mechanism contributes to glial dysfunction in vivo remains undetermined. These results align with earlier studies showing that astrocytes can internalize, process, and release extracellular tau and develop inflammatory responses following tau accumulation^24–26^. In our study, neurons grown with tau-treated astrocytes showed decreased dendritic spine density, branching, and dendritic area. Neutralization of tau, IL-1p, or IL-6 only partly restored dendritic branching. None of the individual neutralization treatments restored dendritic area. This indicates that extracellular tau plays a major role in reduced spine density, while both tau and the cytokines examined contribute to changes in dendritic branching. The continued reduction in dendritic area may reflect other secreted factors, incomplete neutralization, or structural changes not reversed during the experiment. These findings raise the possibility that astrocytes generate and release new, potentially pathogenic tau species following tau uptake and processing, which could contribute to tau propagation^24–26^. Future studies will determine whether these astrocyte-derived tau species contribute to neuronal deficits and whether reducing mitochondrial oxidative stress prevents their generation or release.

hTau mice also exhibited changes in predicted astrocyte-vascular signaling and impaired evoked cerebral blood flow responses (**Figs. 9 and 10**). Transcript-based analyses indicated elevated growth-factor and inflammatory signaling scores between astrocyte and vascular cell populations, suggesting that tauopathy disrupts communication at the gliovascular interface. These interaction scores are based on transcript abundance and do not represent direct measurements of signaling activity. Although astrocyte-targeted SOD2 overexpression altered the cerebral blood flow response profile to more closely resemble wild-type levels, the response area and duration did not differ significantly from those in hTau + GFP mice. Therefore, the SOD2 experiment does not demonstrate a rescue of neurovascular function but supports further investigation into astrocyte mitochondrial oxidative stress as a potential contributor to neurovascular dysfunction. Given that astrocyte mitochondrial reactive oxygen species (ROS) can influence both brain metabolism and vascular function, these findings suggest a possible link between the observed mitochondrial changes and impaired neurovascular coupling^28,32,65^. The in vitro studies utilized a single concentration of tau and only one commercially available preparation of primary human astrocytes, precluding assessment of dose dependence or donor/genotype variability. The cells’ sensitivity to heparin suggests involvement of HSPGs in tau uptake but does not identify the receptor complex or exclude other entry pathways^14,15^. Fluorescence signals from MitoTracker and MitoSOX depend on probe uptake, membrane potential, and oxidation chemistry; however, MitoSOX fluorescence alone does not specifically quantify superoxide levels^66^. Oxygen consumption was not normalized to mitochondrial content, and SOD2 protein abundance was measured without direct assessment of enzyme activity. Additionally, as MitoTEMPO was administered systemically, its effects on astrocyte-enriched fractions cannot be solely attributed to direct action on astrocytes. The in vivo measurements were limited to the cortex, and it remains unknown whether similar changes occur in other brain regions.

In summary, these studies demonstrate that soluble pathogenic tau enters astrocytes and rapidly induces mitochondrial oxidative stress and decreased ATP levels. These early changes occur prior to any detectable microtubule destabilization or induction of senescence-associated markers, and tau exposure also impairs mitochondrial respiration. In hTau mice, MitoTEMPO reduced specific inflammatory and senescence-associated transcripts, supporting the role of mitochondrial oxidative stress in the astrocyte response to tau. Tau-treated astrocytes exerted detrimental effects on neuronal dendritic structure, with extracellular tau, IL-1p, and IL-6 implicated in distinct neuronal outcomes. Collectively, these findings indicate that astrocytes are direct targets of soluble pathogenic tau and support a model in which mitochondrial oxidative stress mediates inflammatory and senescence-associated changes in astrocytes. Whether this pathway directly leads to neuronal injury and neurovascular dysfunction remains to be determined.

## Methods

### Animals

All studies were performed under approval from the Institutional Animal Care and Use Committee, Oklahoma City, Oklahoma, and in accordance with the National Institutes of Health Guide for the Care and Use of Laboratory Animals. The male and female hTau mice, previously described ^67^, produce all six human tau isoforms on a murine Mapt-null background and become susceptible to age-related tau hyperphosphorylation and aggregation; these mice were bred in our animal colony. C57BL/6J mice were used as wild-type (WT) controls. The mice were kept in cages at a maximum density of five per cage, under a 12-hour light and 12-hour dark photoperiod at a temperature of 75 ± 3 °F and with a relative humidity of at least 30% and had free access to food and water. The groups were: 18 months for the single-cell RNA sequencing; 7 and 13 months for the biochemical analysis of acetylated α-tubulin and phosphorylated tau; 7 to 9 months for the measurement of senescence and SASP markers in the whole brain and in the astrocyte-enriched fractions; and 6 months at the time of delivery of the viral vector for the neurovascular coupling study, with functional assessments carried out at 10 months (17 animals in each group). Animals were euthanized by cervical dislocation under ketamine/xylazine anesthesia following assessment of neurovascular coupling, or by isoflurane overdose if not used in vascular measurements.

### Astrocyte isolation from mouse brain and MACS sorting

The cortical tissue was dissociated using the Adult Brain Dissociation Kit, mouse and rat (Miltenyi Biotec, Bergisch Gladbach, Germany; Cat. No. 130-107-677), and debris together with red blood cells were removed as instructed by the manufacturer, after which the astrocytes were positively selected using the Anti-ACSA-2 MicroBead Kit, mouse (Miltenyi Biotec; Cat. No. 130-097-678). The labelled cells were then separated on LS columns (Miltenyi Biotec; Cat. No. 130-041-306) using a MidiMACS Separator (Cat. No. 130-042-302). The columns were washed three times before being removed from the magnet and the cells that had been retained were eluted. The cells were collected by centrifugation and then washed in PBS prior to RNA being collected. The degree of enrichment was assessed by qRT-PCR for astrocyte (Aqp4, Slc1a3, Gfap) versus neuronal (Rbfox3), microglial (Cx3cr1), and endothelial (Pecam1) markers. The isolated fractions were used for qRT-PCR and immunoblot analysis of the senescence and SASP markers.

### Tau uptake

In experiments on tau internalization, primary human astrocytes were seeded on coverslips in 24-well polystyrene plates at 25,000 cells per well and pretreated for 2 hours with either vehicle (PBS) or 1 g/L heparin (Tocris 9041-08-1). The cultures were then exposed to 40 µg/mL recombinant human tau-441, which oligomerized at room temperature in the media for 10 minutes. This tau carried a V5 tag at its C-terminus (Sigma SAE0076). Alternatively, cells were treated with vehicle, with or without heparin, for one hour. The cells were fixed with 4% PFA and permeabilized with Triton X-100. Staining was performed using anti-V5 antibody (Invitrogen R960-25), alpha/beta tubulin (Cell Signaling 2148S), and DAPI. The cells were mounted on slides using ProLong Gold antifade reagent (Invitrogen P36934) and imaged on a Zeiss LSM-710 confocal microscope with a 63x oil objective.

### Primary human astrocyte culture and treatments

The primary human cortical astrocytes were obtained from ScienCell Research Laboratories (Carlsbad, CA, USA; Cat. No. 1800) and were cultivated in 0.1% gelatin-coated cell culture flasks (Sigma-Aldrich; Cat. No. G1890) in a 5% CO_2_-humidified incubator at 37 °C. The culture medium used was DMEM/F-12 (ATCC; Cat. No. 30-2006) to which 10% serum (Cytiva Cosmic Calf Serum) and 0.2% Primocin (InvivoGen; Cat. No. ant-pm-1) were added. The cells, at passage 5, were plated at the same density across all conditions before treatment. The astrocytes were given 40 µg/mL of soluble tau aggregates (O.Tau), an equal amount of KRT8 a protein similar in size and charge to tau or the vehicle (PBS), as mentioned. In order to prevent tau internalization, which is dependent on proteoglycans, the cells were co-treated with 0.5 mg/mL heparin sodium (Tocris)^14^. The treatments were maintained for the durations specified in each figure. For the time-course experiments, cells were collected at 0.5, 1, 2, 3, 4, 6, 8, 12, 24, and 48 hours after treatment, as indicated. For mitochondrial ROS scavenging, astrocytes were pretreated with MitoTEMPO (cayman Item No. 16621) before and during tau exposure^64^. Antimycin A plus rotenone served as a positive control for maximal mitochondrial ROS generation.

### Cell and nuclear size quantification

The human astrocytes were exposed to 0, 100, or 250 µg/mL for 48 hours. Then, the cells were incubated with Hoechst 33342 at 0.3 µg/mL for 30 minutes and imaged with an Operetta High Content Imaging System (Perkin Elmer, Waltham, MA, USA) set at 37 °C and 5% CO2. For each condition, imaging was done in three duplicate wells, with 12 fields examined per well using the Harmony High-Content Imaging analysis platform (Perkin Elmer, Waltham, MA, USA). Nuclear sizes were calculated using cell method A in the Harmony software. All measurements were made while the experimenter was blinded to the treatment groups by randomly selecting 10 to 15 cells in each field for quantitative analysis.

### Mitochondrial ROS, ATP, and membrane potential and mitochondrial protein markers

The level of mitochondrial superoxide was determined using live-cell imaging and the MitoSOX™ Red mitochondrial superoxide indicator (Invitrogen, Thermo Fisher Scientific; Cat. No. M36008). After the experimental treatment, the cells were exposed to 5 µM MitoSOX Red in phenol-free F12 medium (ATCC; Cat. No. 30-2006) that had been prewarmed and kept at 37 °C, protected from light. They were then washed three times with prewarmed medium and imaged immediately using the same imaging settings for each experimental group. The fluorescence from MitoSOX Red was quantified per cell after background subtraction and normalized either to the number of cells or to the nuclear signal detected at 695 nm. An increase in MitoSOX Red fluorescence was taken as evidence of increased mitochondrial superoxide-associated oxidation.

### Capillary electrophoresis

Immunoblotting. In the case of capillary electrophoresis immunoblots, cells and tissue were lysed using 1× Cell Lysis Buffer (Cell Signaling Technology, Cat. No. 9803; prepared from a 10× stock) together with cOmplete™ Mini protease inhibitor cocktail (Roche, Cat. No. 11836153001). The lysates were then examined with the ProteinSimple Wes system. The primary antibodies used were phospho-tau Thr231 (AT180; Thermo Fisher Scientific, Cat. No. MN1040), total tau (Tau5; MilliporeSigma, Cat. No. MAB361), acetylated α-tubulin (clone 6-11B-1; Santa Cruz Biotechnology, Cat. No. sc-23950), α-tubulin (rabbit polyclonal; Cell Signaling Technology, Cat. No. 2144S), and [3-actin (rabbit monoclonal, clone 13E5; Cell Signaling Technology, Cat. No. 4970S). The chemiluminescent signals were quantified as the area under the curve peak using Compass for Simple Western software (ProteinSimple), and the ratio of acetylated α-tubulin to total α-tubulin was calculated as an index of microtubule stability. Antibodies listed for mitochondrial marker analysis were SOD2 (rabbit monoclonal, clone D3X8F; Cell Signaling Technology, Cat. No. 13141), HO-1 (rabbit monoclonal, clone D60G11; Cell Signaling Technology, Cat. No. 5853), and TOMM20 (rabbit monoclonal, clone D8T4N; Cell Signaling Technology, Cat. No. 42406), PGC-1a (rabbit polyclonal; Novus Biologicals, Cat. No. NBP1-04676SS)

### Quantitative real-time PCR

Total RNA from primary human astrocytes and from hTau mouse brain was isolated using the RNAqueous-4PCR kit (Life Technologies), and cDNA was prepared using the Bio-Rad iScript cDNA synthesis kit. Quantitative RT-PCR was performed using the Bio-Rad iTaq Universal SYBR Green Supermix on a 7900HT Sequence Detection System (Applied Biosystems). Ct values were calculated using SDS 2.3 software (Applied Biosystems). Relative mRNA expression was calculated by the comparative method (2“AACT) using GAPDH as a reference^68^. Primers used for qRT-PCR are shown in Supplementary Table 2.

### Single cell RNA sequencing processing and analysis

The cortical single-cell RNA sequencing data from the WT and hTau mice were processed using Cell Ranger to generate filtered feature–barcode matrices. The data were then analyzed with Scanpy^69^ in Python 3.11. Cells with fewer than 200 total counts or fewer than three detected genes were discarded. Quality control was performed separately for each sample. A cell was excluded when the difference between its log-transformed total counts or the number of detected genes and the sample median exceeded 5 median absolute deviations (MADs). Additionally, cells with mitochondrial read percentages more than 3 MADs above the median or higher than 20% were removed. Doublets were detected and eliminated for each sample using Scrublet^70^.

The analysis employed analytic Pearson residuals^71^ and identified 2,000 genes as highly variable using a batch-aware approach. Batch integration was performed using scVI (version 1.4.2 of scvi-tools)^72,73^ with 20 latent dimensions, two layers, up to 100 training epochs, and early stopping. A neighborhood graph was constructed from the scVI latent representation with 30 neighbors. A UMAP embedding was computed with a minimum distance of 0.3. Clustering was performed with Leiden at a resolution of 0.5^74^. Marker genes for the clusters were identified using the Wilcoxon rank-sum test. The cell types were assigned using curated marker panels for the major CNS and vascular populations. Clusters dominated by hemoglobin, platelet, or ambient RNA-associated signatures, and those with low-complexity profiles, were excluded as technical artifacts. The astrocytes were divided for further analysis. The genes with the highest variability were selected again for further analysis. A separate scVI model was trained with 15 latent dimensions, two layers, and 80 epochs. Astrocyte substates were determined by applying Leiden clustering at a resolution of 0.4. Differential expression was evaluated using the Wilcoxon rank-sum test, with multiple testing correction using the Benjamini–Hochberg method^75^; genes with adjusted P-values < 0.05 were considered significant. Genes associated with sex chromosomes, such as Xist, Tsix, Ddx3y, Eif2s3y, Uty, and Kdm5d, were excluded. The treatment-matched comparison shown in **Fig. 3** included two WT and two vehicle-treated hTau animals. The genes were ordered by their signed Wilcoxon test statistic as part of a preranked gene set enrichment analysis using 1,000 permutations^76^. The gene sets were obtained from the MSigDB Hallmark (2020)^77^, Gene Ontology Biological Process (2023), and KEGG (2019, mouse) collections. Normalized enrichment scores and false discovery rate q-values were provided. The leading-edge genes that contributed to the enrichment signals that were selected were extracted, and the per-cell scores were calculated by means of Scanpy’s score_genes function.

For the treatment-matched group consisting of two wild-type and two hTau animals, summaries of gene expression levels were prepared. Log-normalized expression values were averaged per animal, then transformed into z-scores for each gene. Group comparisons were performed using the Wilcoxon rank-sum test, with each animal serving as a biological replicate. The analyses were considered exploratory due to the small sample size. Intercellular signaling was examined using a CellPhoneDB-style permutation framework^78^, with 1,000 permutations performed separately for each genotype. Analyses focused on astrocytes and on various vascular-associated populations, such as endothelial cells, a second endothelial-like cluster, pericytes/mural cells, and vascular leptomeningeal cells/fibroblasts. The second endothelial-like cluster was retained, but with lower confidence in its annotation; canonical endothelial markers such as Pecam1, Cldn5, Flt1, and Slco1a4 were found in more than 85% of cells in this cluster.

For each directional source–target pair, the interactions that reached an uncorrected permutation threshold of P < 0.01 were counted. Those interactions that met the threshold only in the hTau group or only in the WT group were referred to as gained in hTau or lost in hTau, respectively. These designations reflect genotype-specific threshold membership rather than on direct statistical tests comparing the two genotypes. The selected interactions were shown with their permutation P-values and interaction strength values based on mean expression.

The interactions were grouped into functional categories based on literature-derived gene annotations, including growth-factor signaling, Wnt/vascular stability, extracellular matrix/adhesion, chemokine/inflammatory signaling, and cell guidance/trophic signaling. This classification was merely descriptive and did not constitute formal gene set enrichment.

### Primary neuron culture and astrocyte-neuron coculture

Primary mouse neurons were obtained from embryos on embryonic day 18 (E18) from CD1 female mice using a previously described prenatal-neuron isolation protocol^79^. They were plated at 100,000 cells per well onto poly-D-lysine-coated sterile glass coverslips in 24-well plates. The neurons were grown in Neurobasal medium (Gibco, Cat. No. 21103-049) with 1 mM GlutaMAX (Gibco), 2% B27, and 1% penicillin-streptomycin (Sigma) at 37 °C in a humidified 5% CO_2_ incubator. The plating medium also contained 2% fetal bovine serum and was replaced with standard neuronal medium after the cells attached (3-6 h). Primary human astrocytes were plated at 10,000 cells per well in 0.4-µm-pore polycarbonate transwell inserts (Thermo Fisher Scientific; Cat. No. 140620). They were treated with 40 µg/mL of soluble tau aggregates or vehicle for 48 h, then washed three times with Hank’s balanced salt solution on both inner and outer surfaces of each insert. The inserts were transferred into wells containing neurons at 18 days in vitro. Neurobasal medium was used in both chambers, and co-cultures were kept for 96 h with one medium change on day 3. For the neuron-only control, inserts without astrocytes were used.

In the antibody-neutralization experiments, 500 ng/mL of nonspecific IgG (mouse; Abcam; Cat. No. ab188776), anti-IL-1p (Abcam; Cat. No. ab2105), anti-IL-6 (rabbit polyclonal; Abcam; Cat.No. ab6672), or anti-tau (mouse monoclonal, clone Tau5; Millipore; Cat. No. MAB361) was added to the lower chamber throughout the 96-hour coculture. IgG was used as a control for nonspecific antibodies. Although the experiments aimed to assess the contributions of IL-1p, IL-6, and tau to the neuronal effects caused by astrocytes exposed to tau, they do not exclude the possibility that other secreted factors also contributed.

The neurons were fixed at room temperature for 30 minutes in 4% paraformaldehyde, treated with 0.2% Triton X-100 to permeabilize them, and then stored at 4 °C overnight with the anti-MAP2 antibody (mouse monoclonal, clone HM-2; Sigma; Cat. No. M4403). After washing, the cells were incubated with Alexa Fluor 594-conjugated goat anti-mouse secondary antibody (Life Technologies; Cat. No. A11032) and stained with DAPI. Z-series images were obtained using a Zeiss LSM 780 confocal microscope and a 64× oil-immersion objective. The dendrites were reconstructed and spine morphology analyzed using Imaris FilamentTracer in manual mode (Bitplane, Zurich, Switzerland).

### Astrocyte-targeted SOD2 overexpression

Six-month-old hTau mice received a single retro-orbital injection of AAV-PHP.eB a capsid engineered for efficient, noninvasive CNS transduction following intravenous administration^58^, with the injection containing either GFAP-hSOD2-P2A-GFP or GFAP-GFP as a control, at 1 × 10¹¹ vector genomes per mouse. The compact GfaABC1D promoter was employed in order to limit the expression to astrocytes^59^. SOD2 is a superoxide scavenger in the mitochondrial matrix; its overexpression reduces hippocampal superoxide and prevents memory deficits in AD model mice^62^, and astrocyte-specific SOD2 enhancement mitigates senescence and cognitive impairment associated with aging^80^. The efficient noninvasive transduction of the CNS by AAV-PHP.eB after systemic delivery is largely due to the Ly6a allele found in C57BL/6J mice and this effect is considerably diminished in other genetic backgrounds^81,82^; we did not independently benchmark transduction efficiency in our hTau/Mapt-null background against a standard C57BL/6J cohort. Expression was allowed to proceed for 4 months, and neurovascular coupling was assessed at 10 months of age (n = 17/group).

### MitoTEMPO injection in hTau mice

hTau mice received MitoTEMPO or vehicle at 1.5 mg/kg, administered intraperitoneally once daily for two weeks, after which brains were collected for astrocyte isolation and magnetic-activated cell sorting as described above (**Fig. 7A**).

### Evoked Neurovascular coupling measurements

The cerebral blood flow (CBF) responses were obtained by means of laser Doppler flowmetry. The mice were anaesthetized with ketamine (85 mg/kg) and xylazine (15 mg/kg), intubated, mechanically ventilated and then positioned in a stereotaxic frame on a heating pad to keep their body temperature constant. During the entire procedure, blood pressure was measured using a CODA monitor (Kent Scientific), and the heart rate, peripheral oxygen saturation and end-tidal CO_2_ were continuously monitored with a PhysioSuite system (Kent Scientific). The bone covering the left somatosensory barrel cortex (this area being situated 0.5 to 1.5 mm posterior and 3 to 4.5 mm lateral to the bregma) was removed, a silicone barrier was prepared around the craniotomy, artificial cerebrospinal fluid (aCSF) was directed over the exposed cortex and a Transonic laser Doppler probe was put onto the barrel cortex. After a stable baseline had been achieved, functional hyperaemia was induced by giving contralateral electrical stimulation to the whisker pad at 5 Hz and 1 mA for 30 seconds. For each animal, five stimulation trials were performed and then averaged. The responses were recorded under three consecutive superfusion conditions: (i) aCSF alone; (ii) the selective neuronal nitric oxide synthase inhibitor N^A^co-propyl-L-arginine (L-NPA, 200 nM); and (iii) L-NPA together with the nonselective nitric oxide synthase inhibitor N^A^co-nitro-L-arginine methyl ester (L-NAME, 10 pM). This sequence of pharmacological treatments was used in order to identify the nNOS-dependent, the remaining L-NAME-sensitive and the non-NOS-dependent components of the evoked CBF response described here^83^.

### Statistical analyses

The statistical analyses were performed using GraphPad Prism (La Jolla, CA, USA). In order to compare the means of two experimental groups, a two-tailed unpaired t-test was used. When the variances were found to be unequal, a t-test including Welch’s correction was applied. One-way ANOVA was carried out when appropriate, followed by Tukey’s or Šídák’s post hoc tests. Homogeneity of variance was evaluated using the Brown-Forsythe test; if the variances were unequal, a Kruskal-Wallis test was conducted with Dunn’s multiple-comparisons test, and a Bonferroni correction was applied. These corrections were applied within each group comparison for a particular marker; a p-value less than 0.05 was considered significant. The Grubbs’ test was employed to identify statistical outliers (with α = 0.05). Unless otherwise stated, n refers to the number of independent biological replicates (that is, individual animals or independently derived and treated astrocyte preparations), not the number of technical replicates. The sample sizes were decided upon on the basis of the effect sizes and the variability observed in pilot experiments and in previous work with this system, rather than as a result of a prior power calculation. Both male and female mice were included in all the in vivo experiments; since the study was not designed to detect sex differences, the data from both sexes were combined for analysis.

## Acknowledgments

We acknowledge the following funding support: NIH National Institute on Aging (NIA) 1R01AG057964-01 (Galvan) and US Department of Veterans Affairs I01 BX002211-01A2 (Galvan), the Oklahoma Nathan Shock Center on Aging (Galvan) NIA 2-P30AG050911-11, San Antonio Medical Foundation (Galvan), the JMR Barker Foundation (Galvan), and a William & Ella Owens Medical Research Foundation Grant. Dr. Galvan is the recipient of a Research Career Scientist award, IK6 BX006026, from the US Department of Veterans Affairs. We acknowledge the US Department of Veterans Affairs Biomedical Laboratory Research and Development Service IK2 BX003798-01A1 and a pilot award from the Oklahoma City Veterans Health Care System (Hussong). Dr. Galvan acknowledges the generous support from the Robert L. Bailey and daughter Lisa K. Bailey Alzheimer’s Fund in memory of Jo Nell Bailey. These studies used the Healthspan and Functional Assessment Core and the Pathology Core of the San Antonio Nathan Shock Center of Excellence in the Biology of Aging (NIH/NIA 2 P30 AG013319-21). Some images were generated in the Core Optical Imaging Facility, which is supported by UTHSCSA, NIH-NCI P30 CA54174 (CTRC at UTHSCSA) and NIH-NIA P01A; and the San Antonio Nathan Shock Center Pathology Core (NIH/NIA 2 P30 AG013319-21).

